# Native RNA sequencing on nanopore arrays redefines the transcriptional complexity of a viral pathogen

**DOI:** 10.1101/373522

**Authors:** Daniel P. Depledge, Kalanghad Puthankalam Srinivas, Tomohiko Sadaoka, Devin Bready, Yasuko Mori, Dimitris G. Placantonakis, Ian Mohr, Angus C Wilson

**Affiliations:** Department of Microbiology, New York University School of Medicine, New York, NY 10016, USA; Division of Clinical Virology, Center for Infectious Diseases, Kobe University Graduate School of Medicine, 7-5-1 Kusunoki-cho, Chuo-ku, Kobe, 650-0017, Japan; Department of Neurosurgery, New York University School of Medicine, New York, NY 10016, USA; Kimmel Center for Stem Cell Biology, New York University School of Medicine, New York, NY 10016, USA; Laura and Isaac Perlmutter Cancer Center, New York University School of Medicine, New York, NY 10016, USA; Brain Tumor Center, New York University School of Medicine, New York, NY 10016, USA; Neuroscience Institute, New York University School of Medicine, New York, NY 10016, USA

**Keywords:** Nanopore sequencing, minION, native RNA sequencing, transcriptomics, herpes simplex virus, human herpesvirus 1

## Abstract

Viral genomes exhibit a higher gene density and more diversified transcriptome than the host cell. Coding potential is maximized through the use of multiple reading frames, placement of genes on opposing strands, inefficient or modified use of termination signals, and the deployment of complex alternative splicing patterns. As a consequence, detailed characterization of viral transcriptomes by conventional methods can be challenging. Full length native RNA sequencing (nRNA-seq) using nanopore arrays offers an exciting alternative. Individual transcripts are sequenced directly, without the biases inherent to the recoding or amplification steps included in other sequencing methodologies. nRNA-seq simplifies the detection of variation brought about by RNA splicing, use of alternative transcription initiation and termination sites, and other RNA modifications. Here we use nRNA-seq to profile the herpes simplex virus type 1 transcriptome during early and late stages of productive infection of primary cells. We demonstrate the effectiveness of the approach and identify a novel class of intergenic transcripts, including an mRNA that accumulates late in infection that codes for a novel fusion of the viral E3 ubiquitin ligase ICP0 and viral membrane glycoprotein L.

## INTRODUCTION

Herpesviruses are adept viral pathogens that have co-evolved with their hosts over millions of years. Like all viruses their success is predicated on repurposing of the host transcriptional and translational machinery^1, 2^ and through use of compact, gene-dense genomes with exceptional coding potential^3–7^. The 152 kb double-stranded DNA genome of herpes simplex virus type 1 (HSV-1) includes roughly 80 genes, predominantly single exon open reading frames (ORFs), some transcribed as polycistronic mRNAs, along with a smaller number non-coding RNAs^8, 9^. These are traditionally grouped into three kinetic classes^10–12^. Although splicing of HSV-1 mRNAs is infrequent, exceptions include mRNAs encoding ICP0, ICP22, UL15p, and ICP47, as well as the non-coding latency-associated transcripts.

Conventional RNA sequencing methodologies, while highly reproducible, utilize multiple recoding steps (e.g. reverse transcription, second strand synthesis and in some cases, PCR amplification) during library preparation that may introduce errors or bias in the resulting sequence data^13^. Data quality may be further convoluted by the use of short-read sequencing technologies, which require well-curated reference genomes to accurately assess the abundance and complexity of transcription in a given system. Loss of information on transcript isoform diversity including splice variants, is especially problematic^14^. Despite these inherent difficulties, recent studies have shown that host transcription and mRNA processing are extensively remodeled during HSV-1 infection^15–17^ and recent studies using cDNA-based short- and long-read sequencing technologies indicate that the HSV-1 transcriptome is substantially more complex than previously recognized^18–20^.

To examine this in more detail, we have employed a new methodology for single molecule sequencing of native mRNAs (nRNA-seq) using nanopore arrays^21^. Specifically, we have used the Oxford Nanopore Technologies MinION platform to directly sequence host and viral mRNA from infected human primary fibroblasts at early and late stages of infection. Error correction, a pre-requisite for current nanopore sequence read datasets, and the generation of pseudo-transcripts were guided using Illumina short-read sequence data from the same source material.

In addition to highlighting the fidelity and reproducibility of native mRNA nanopore sequencing, we also provide a new approach to error correction that results in error-free transcript sequences from which internal ORFs can be accurately translated to predict protein sequences. Finally, we describe a series of important observations about the HSV-1 transcriptome, including the existence of multiple new transcription initiation sites that produce mRNAs encoding novel or alternative ORFs, and the existence of spliced HSV-1 read through transcripts that encode novel protein fusions and provide evidence for disruption of transcription termination for a number of viral transcription units. We show that one of these, a fusion between the ORFs encoding the viral E3 ubiquitin ligase ICP0 and viral membrane glycoprotein L, produces a 32-kDa polypeptide expressed with late kinetics. Taken together, these studies demonstrate the ability of nRNA-seq to identify novel mRNA isoforms that further expand the coding potential of gene dense viral genomes.

## RESULTS

### Combined sequencing of host and viral transcriptome using nanopores

To evaluate the reproducibility of native RNA sequencing (nRNA-seq) using nanopore arrays we prepared total RNA from two biological replicates of normal human dermal fibroblasts (NHDF) infected with HSV-1 GFP-Us11 strain Patton (hereafter HSV-1 Patton)^22, 23^ for 18 hours (Fig. 1a, Supplementary Fig. 1a, Supplementary Table 1). Sequencing libraries were generated from the poly(A)+ RNA fraction and sequenced on a MinION MkIb with R9.4 flow cell with a run time of 18 h, yielding up to ~550,000 reads (Fig. 1a, Supplementary Fig. 1a) that were then aligned against the human transcriptome and HSV-1 strain 17 *syn*^+^ annotated reference sequence using the splice-aware aligner MiniMap2^24^. Relative transcript abundances for both host and viral mRNAs showed high reproducibility between biological replicates (*H. sapiens* r^2^ = 0.985, HSV-1 r^2^ = 0.999, (Fig. 1b) and minimal mRNA decay during library construction and sequencing (Supplementary Fig. 1b). Next, we compared the 18 h nanopore sequence data to an Illumina mRNA-Seq dataset generated from the same source RNA (Fig. 1c). Here, the reverse transcription and amplification steps utilized in standard Illumina protocols, combined with the reduced accuracy inherent in mapping short read data, resulted in similar but not identical profiles, as measured either by relative viral transcript abundances or by a genome-wide sliding window approach (Supplementary Fig. 1c). As a final examination, we constructed an additional nRNA-seq library from the same source material (technical replicate) and ran this on a separate MinION device, confirming that the sequencing data were also reproducible across instruments (Supplementary Fig. 1d). Satisfied that nanopore nRNA-seq is highly reproducible, we subsequently sequenced two additional samples to enable comparisons between early (6 h) and late (18 h) time points of HSV-1 Patton infection of NHDFs, and to examine the contribution of the virion host shut off (vhs) protein (Fig. 1a, Supplementary Fig. 1a, Supplementary Table 1). Both of these yielded similar sized datasets with the major difference being a significantly reduced fraction of HSV-1 sequence reads in the Δvhs dataset (Supplementary Table 1), likely reflecting the involvement of vhs in antagonizing innate defenses at the beginning of the infection cycle^25^.

**Fig. 1:**
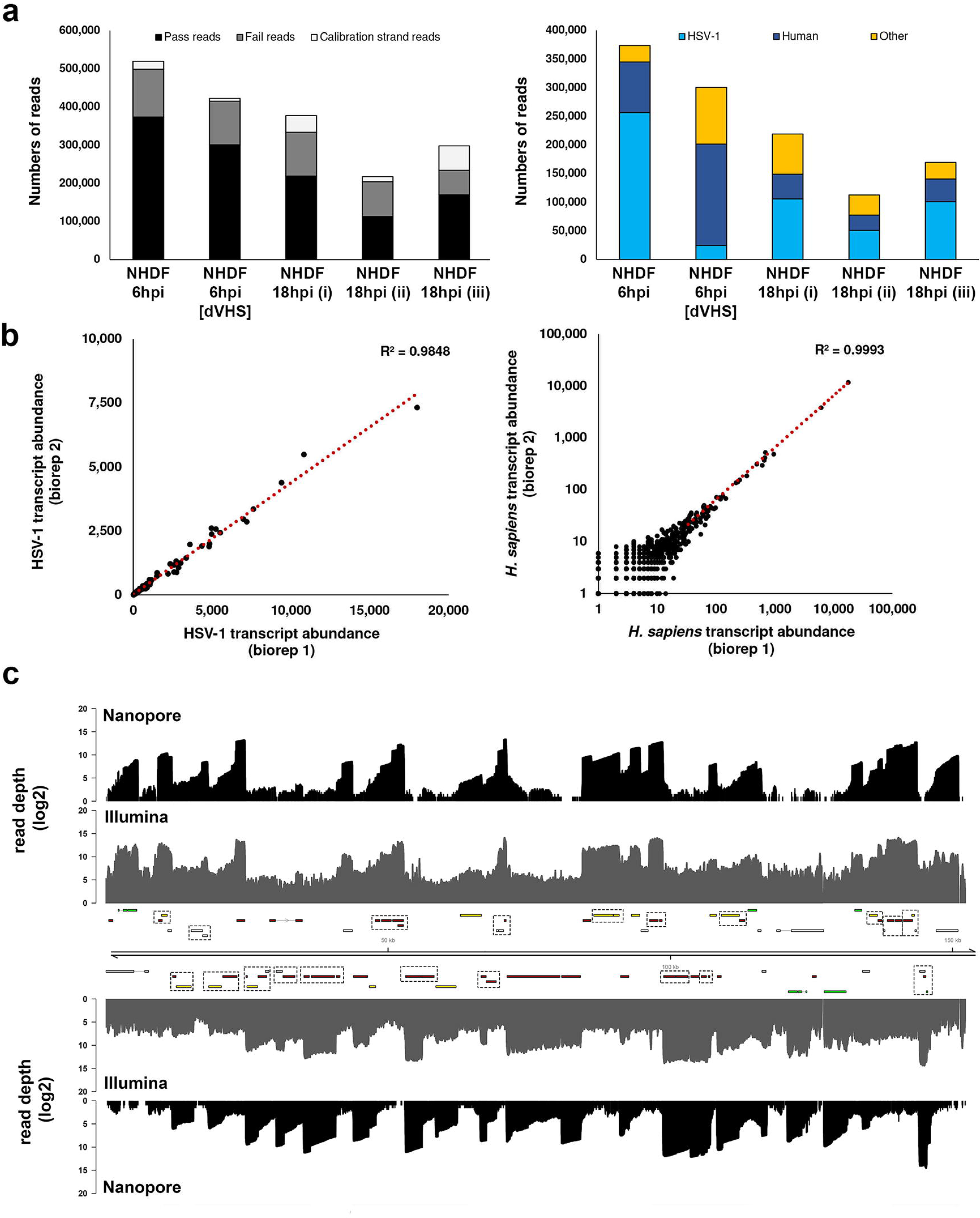
Native RNA sequencing using nanopore arrays is highly reproducible. **a**, Summary metrics for five separate nRNA-Seq runs using normal human dermal fibroblasts (NHDFs) infected with HSV-1 strain Patton GFP-Us11 or HSV-1 strain F vhs null (dVHS) for either 6 or 18 h. NHDF 18hpi (i) and (ii) represent biological replicates, with an additional technical replicate, NHDF 18hpi (iii), performed on a separate minION device. Calibration strand reads originate from the spiked human enolase 2 (ENO2) mRNA. Pass and fail reads were classified as such by the albacore basecaller. Only reads passing QC were utilised for analysis and mapped against the HSV-1 and *H. sapiens* transcriptomes. Only a small proportion of reads could not be mapped. **b**, transcript abundances were counted for the HSV-1 (left) and *H. sapiens* (right) transcriptomes and showed near perfect correlation between biological replicates of the NHDF 18hpi (i) and NHDF 18hpi (ii) samples. **c**, HSV-1 genome wide coverage plots of poly(A) RNA sequenced by nanopore (black) and Illumina (dark grey) technologies. Note that nanopore reads represent a single mRNA, directly sequenced, while Illumina reads are derived from highly fragmented mRNAs that have undergone reverse transcription and multiple round of amplification. The HSV-1 genome is annotated with all canonical open reading frames (ORFs) and coloured according to kinetic class (green - immediate early, yellow - early, red - late, grey - undefined). Multiple ORFs are grouped in polycistronic units and these are indicated by hatched boxes.

### The utility of error-correction and pseudo-transcripts

Each nanopore-derived sequence read represents a single full length or 5’ truncated transcript present in the starting poly(A)+ RNA pool and maps with high specificity but low identity (80-90%) to the reference genome/transcriptome. As polyadenylated transcripts are likely translated within the infected cell, we asked if we could identify known or novel open reading frames (ORFs) within our sequence reads and thereby determine the breadth of protein variants expressed by the virus in these cells. This approach was hindered by the presence of numerous indels and substitution type errors within the raw nRNA-seq reads, a consistent issue with nanopore sequencing. To overcome this, we designed a novel error-correction strategy utilizing proovread^26^ to correct the raw sequence read data (Fig. 2a), combined with a decision matrix operating across a range of error-corrected subsampled datasets (Fig. 2b). Briefly, we assessed the amount of error in a given read by determining the CIGAR string length for that read in the uncorrected dataset versus corrected datasets generated using Illumina RNA-Seq data from the same source material (Fig. 2a). The optimal version of an error-corrected read was considered to have the shortest CIGAR string length amongst all subsampled datasets (Fig. 2b). Our approach showed that error-correction notably reduces the numbers of indel and substitution-type errors in all nanopore reads (Supplementary Fig. 2), the impact of Illumina read subsampling was minimally affected by the size of the subsampled dataset (Supplementary Fig. 2), error correction rescued up to 9% of reads that could not be mapped previously (Fig. 2c). Despite the relative success of error-correction, sufficient indel errors remained to preclude accurate identification of encoded ORFs. We therefore used the mapping positions resulting from alignment of the error-corrected nanopore reads to generate ‘pseudo-transcripts’. Each pseudo-transcript contains the same alignment start, stop and internal splicing sites as the error-corrected read but with the transcript sequences corrected by substituting in the corresponding reference genome sequence (Fig. 2a). For each pseudo-transcript, we generated translations of all encoded ORFs greater than 90 nucleotides in length and confirmed the presence of full length mRNAs encoding the expected protein products for all canonical HSV-1 genes except UL36 and UL52, two of the longest templates. While pseudo-transcripts could theoretically be generated from mapping raw nanopore reads, error-correction significantly improves the accurate prediction of splice sites – a crucial factor when attempting to identify novel transcript isoforms.

**Fig. 2:**
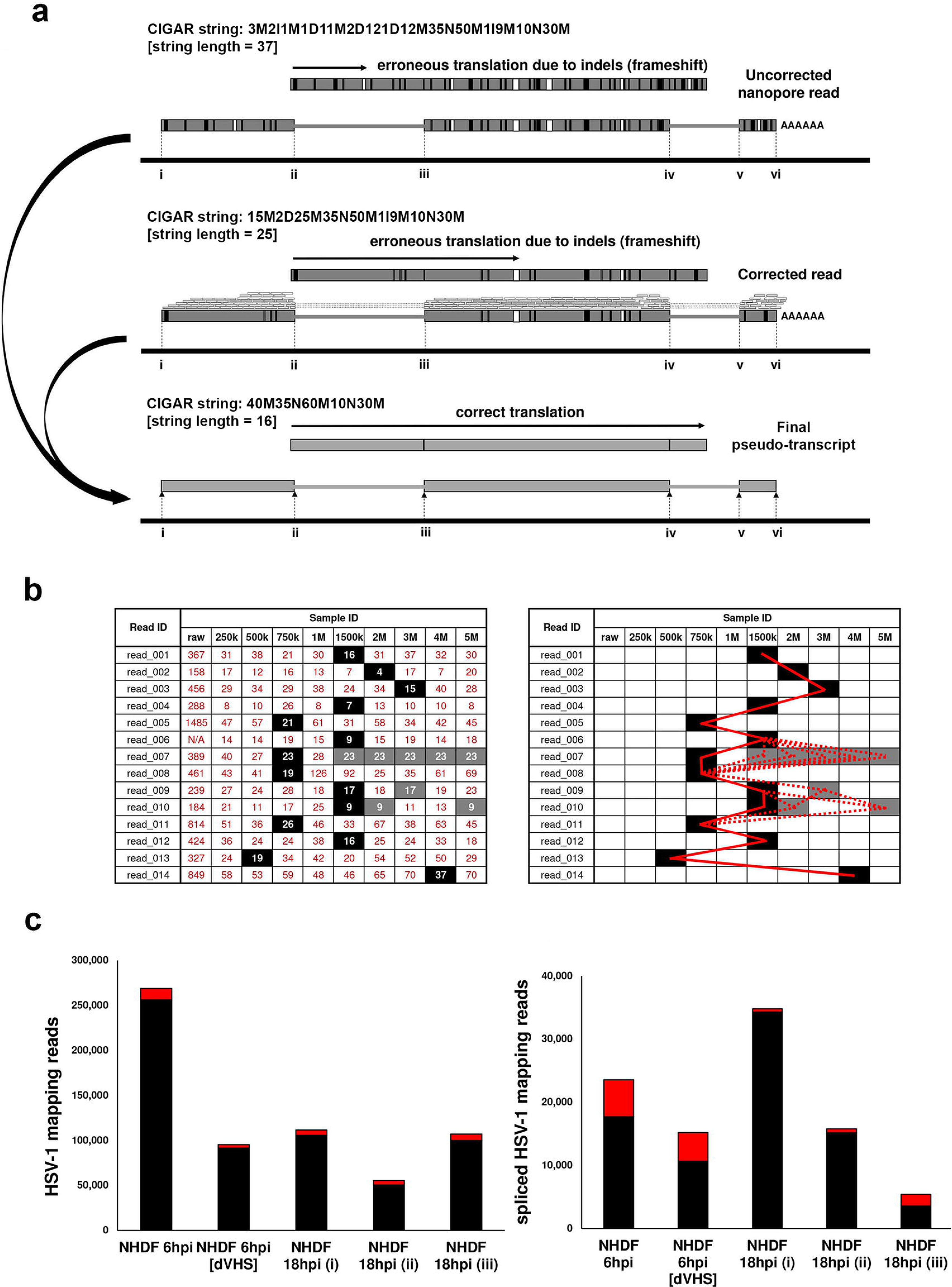
Error-correction and generation of pseudo-transcripts to overcome sequencing errors inherent to the nanopore method. **a**, Raw direct RNA nanopore reads include numerous indel and substitution errors that hinder the identification of encoded ORFs and thereby impede annotation of the transcriptome. Illumina datasets generated from the same material allowed error-correction using proovread (and see Figure S2). Subsequently the transcript start / stop positions and internal splice positions were used to generate pseudo-transcripts free of indel and substitution errors that permit unambiguous ORF prediction. Example changes in CIGAR string lengths for a given read are shown for each step of correction. **b**, to optimise proovread error-correction, we tested a range of subsampled Illumina datasets and evaluated corrected reads by the length of the CIGAR string (see methods). Because optimal Illumina subsampling varies between reads, we subsequently applied a decision matrix utilising the best-corrected version of a given read (filled boxes) as scored by smallest CIGAR string length. Where multiple subsampling sets produce identical shortest CIGAR scores (shaded boxes), no difference was observed between resulting sequences. The bold red line indicates the path chosen (i.e. from which error-corrected dataset a given read was drawn) while the dotted lines indicate alternative paths that produce the exact same result due to having identical CIGAR string lengths. **c**, error correction rescued a proportion of unmapped reads and reassigned these as HSV-1 mapping (left) and improved splice site detection in all datasets (right). Black bars indicate the HSV-1 mapping reads (left) and spliced HSV-1 reads (right) in the uncorrected dataset. Red bars indicate the numbers of rescued reads following error correction.

### Profiling viral transcription at early and late stages of infection

A primary goal of our study was to evaluate the coding capacity of HSV-1 in the context of alternative and novel transcription. The distribution of nRNA-seq read lengths for both viral and host transcripts remained similar at discrete sampling times (6 and 18 h) and interestingly, were not obviously different for the HSV-1 Δvhs mutant (data not shown). It is possible the vhs endonuclease has limited impact on the viral transcriptome at the times examined, or that once cleaved, mRNAs are degraded very rapidly. Peaks representing the 5’ and 3’ ends of sequenced mRNAs map closely to previously established transcription start and termination sites of many HSV-1 transcription units and we term the peak locations as proximal transcription start sites (pTSS) and transcription termination sites (pTTS). We noted canonical TATAAA boxes upstream of thirteen HSV-1 genes, with the maximal pTSS peak positioned 30 to 48 bp downstream. We catalogued proximal pTSS sites for all canonical HSV-1 ORFs except for UL9, UL36, UL52, and RS1, as well as the latency-associated transcript precursor (Supplementary Table 2). While use of these pTSS was consistent between the 6 and 18 h time points, several ORFs (UL6, UL8, UL14, UL24, UL29, UL51, UL53, US1) were notable for the apparent use of multiple pTSS (Fig. 3, Supplementary Fig. 3 and 4, Supplementary Table 3) resulting in elongated 5’ UTRs, or 5’ truncated transcripts. Moreover, we observed eight pTSS located within previously defined ORFs (UL6, UL12, UL24, UL30, UL41, UL44, UL53, UL54, see Supplementary Table 4), potentially encoding novel (different reading frame) or alternative (same reading frame) protein products or polyadenylated non-coding RNAs (ncRNAs). By contrast, cataloguing the pTTS sites revealed that the sites of cleavage and polyadenylation remained relatively constant for viral transcripts, terminating close to canonical A[A/U]UAAA poly(A) signal sequences (Supplementary Fig. 3).

**Fig. 3:**
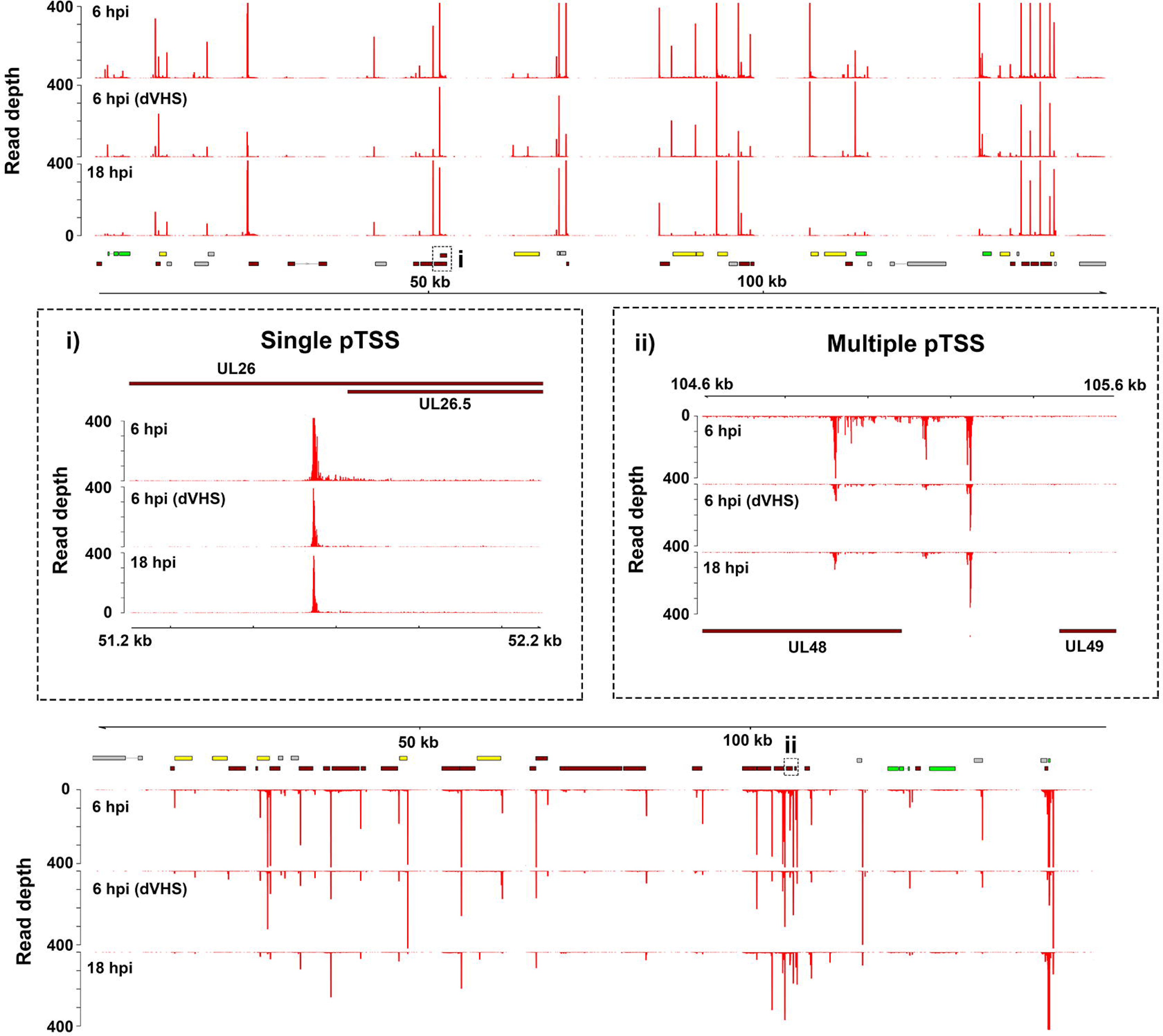
Viral mRNAs initiate at single or multiple locations. To visualize transcription start site positions, the extreme 5’ end of each nRNA-seq read was plotted against the HSV-1 genome. Data sets correspond to NHDF infected with HSV-1 strain Patton for 6 hpi (upper track), strain F dVHS for 6 hpi (middle track) and strain Patton for 18 hpi (lower track). Peaks corresponding to clustered 5’ ends, are referred to as proximal transcription start sites (pTSS) and likely differ by only a few nucleotides from the actual capped mRNA 5’ end. In thirteen cases pTSSs are positioned 30-48 bp downstream of canonical TATA boxes. Upper panel: top strand. Lower panel: bottom strand. Inset boxes: (i) Transcription of the HSV-1 UL26.5 gene initiates at a single location throughout infection. (ii) UL48 transcription initiates at multiple locations, one of which is internal to the canonical UL48 ORF, suggesting transcripts encoding a truncated or alternative protein. Canonical HSV-1 ORFs are colored according to kinetic class (IE – green, E – yellow, L – red, undefined – grey) while polycistronic transcriptional units are indicated by hatched boxes.

### Fusion transcripts further increase the coding capacity of viral genomes

Visual inspection of read alignments using the integrative genomics viewer (IGV)^27^ provided evidence of read-through transcription at both 6 h and 18 h, meaning transcript sequences that contained canonical poly(A) signal sequences at least 500 nucleotides upstream of the actual 3’ transcript end. While read-through transcription across the host genome can leads to aberrant non-adenylated transcripts^18^, here, the compact nature of the HSV-1 genome enables transcription to correctly terminate at poly(A) signal sites further downstream, often at the end of adjacent single or polycistronic transcription units. HSV-1 transcripts with internalized poly(A) signal sites at least 500 nucleotides upstream of the observed 3’ end accounted for around 8% of all HSV-1 nRNA-seq reads in each dataset. In other words, the frequency of read-through transcription was almost the same at 6 and 8 h of infection (Fig. 4a). Read-through transcription provides a simple mechanism for generation of chimeric polyadenylated RNAs. Chimeric RNAs in mammalian cells are thought to arise predominantly from *trans*-splicing^28^ but in compact, gene-dense viral genomes, read through could produce multi-ORF transcripts in which *cis*-splicing^29^ between neighboring ORFs creates novel fusion proteins (Fig. 4b). To search for examples, we predicted the translation products of all spliced HSV-1 transcripts and compared the results to a database of all canonical ORFs. This revealed three distinct fusions of neighboring ORFs (RL2-UL1, UL52-UL54, and US1-US3), (Fig. 4c, Fig. 5, and Supplementary Table 5). The canonical splice donor and acceptor motifs used in each of these fusions were perfectly conserved across more than 150 HSV-1 genomes suggesting the new products are functional (Supplementary Table 5).

**Fig. 4:**
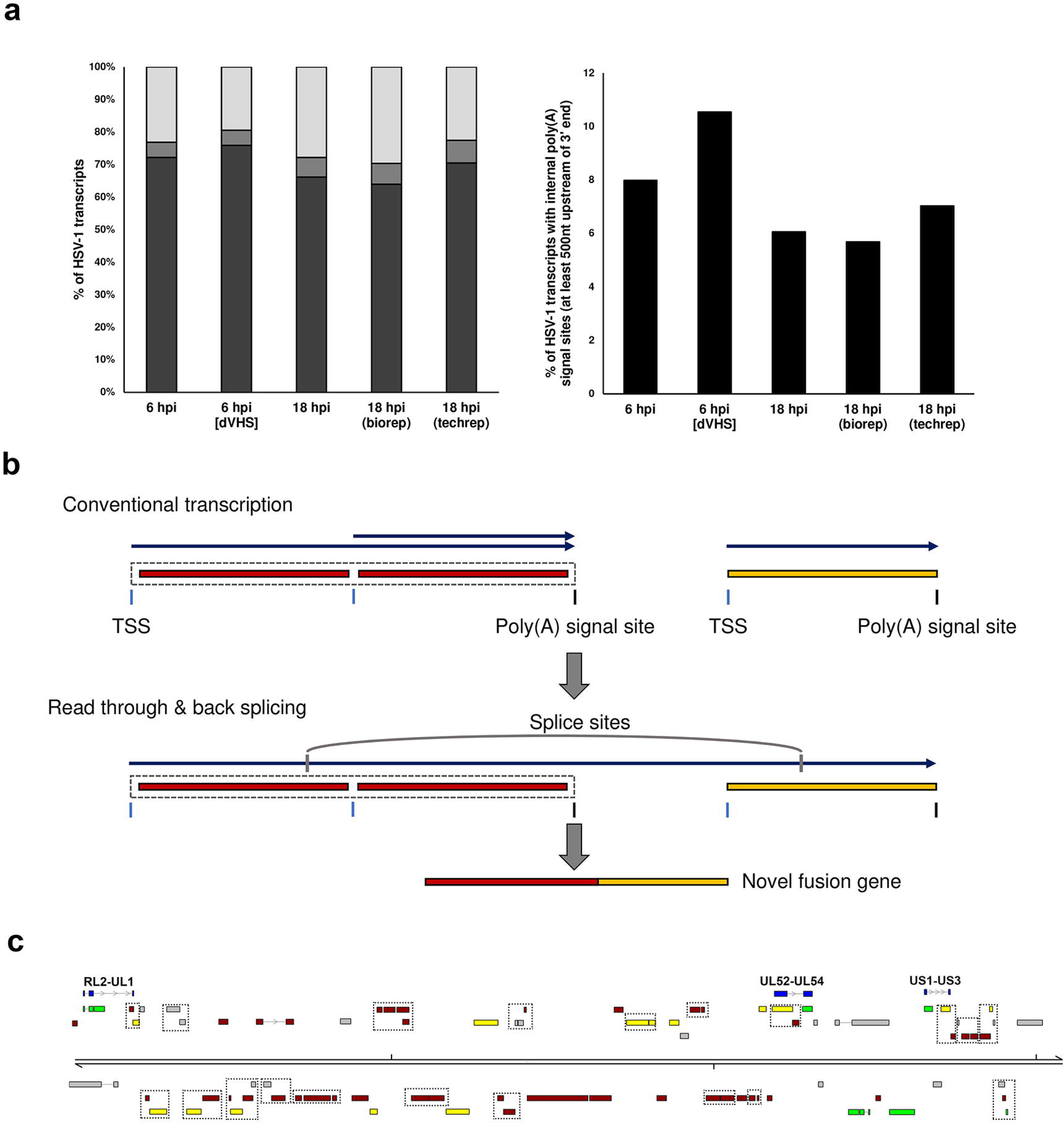
Read through transcription and splicing give rise to multiple HSV-1 fusion transcripts. **a**, (left) HSV-1 transcription termination is predominantly initiated by recognition of a canonical (AAUAAA - dark grey) poly(A) signal sequence although a common variant (AUUAAA - medium grey) and other less well-defined sequences may also contribute (light grey). (right) HSV-1 transcripts with internalized poly(A) signal sites (> 500 nt upstream of transcription termination) accounted for up to 11% of all HSV-1 transcripts at early and late stages of infection. **b**, Transcription of HSV-1 genes initiates at transcription start sites (TSS, blue vertical line) and typically terminates shortly after traversing a canonical (A(A/U)UAAA) poly(A) signal sites (black vertical line). In rarer cases, termination does not occur and transcription extends further downstream as read through until another poly(A) signal site is used. These extended transcripts may be subject to internal splicing which can give rise to fusion ORFs. **c**, schematic representation of the novel fusion ORFs RL2-UL1, UL52-UL54, and US1-US3 (indicated in blue). Canonical HSV-1 ORFs are colored according to kinetic class (IE – green, E – yellow, L – red, undefined – grey) while polycistronic transcriptional units are indicated by hatched boxes.

**Fig. 5:**
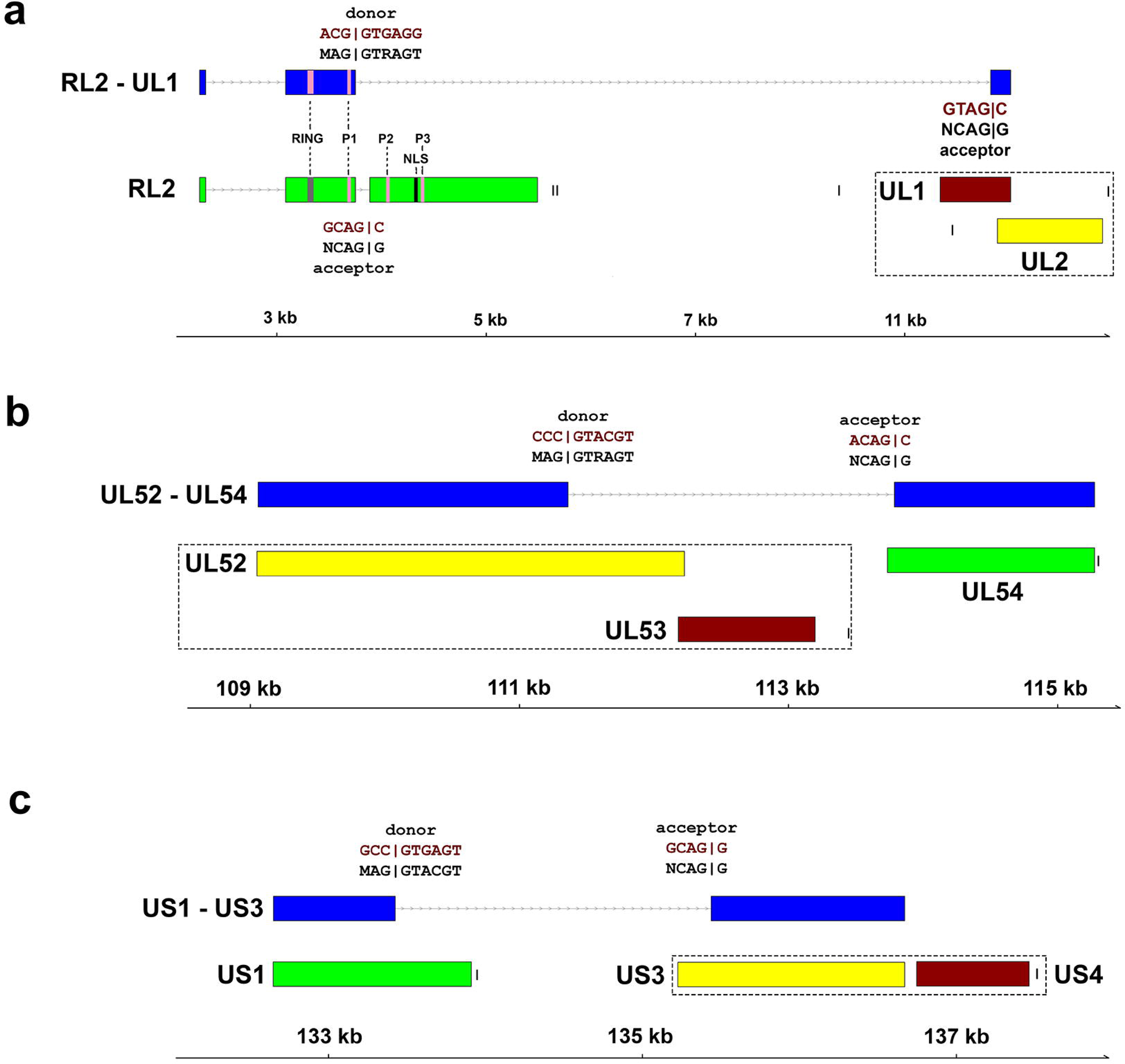
Schematic representation of three novel fusion ORFs. Fusion ORFs are predicted to arise from read through transcription followed by back splicing. Here, all three fusion ORFs arise due to splicing using near-canonical splice donor and acceptor sequences that are perfectly conserved across available HSV-1 genome sequences. **a**, the RL2-UL1 fusion transcript encodes an ICP0-gL fusion protein that lacks two phosphorylation sites (P2, P3) and the nuclear localization signal (NLS) domain present in ICP0. **b**, the UL52-UL54 fusion. **c**, the US1-US3 fusion. Canonical HSV-1 ORFs are labelled and colored according to kinetic class (IE – green, E – yellow, L – red, undefined – grey) while polycistronic transcriptional units are indicated by hatched boxes. Canonical (AAUAAA) poly(A) signal sequence are indicated by black vertical lines. Canonical splice donor and acceptor sequences are shown in black text, with the actual sequences in red text.

### Chimeric RL2-UL1 mRNA is expressed with late kinetics and encodes a novel ICP0 and glycoprotein L fusion

The in-frame fusion of ICP0 (ORF RL2) and glycoprotein L [gL] (ORF UL1), arises through splicing from the canonical exon 2 splice donor within the coding sequence of RL2 to a novel splice acceptor with the UL1 ORF, such that the first two exons of RL2 (residues 1 to 241) including the RING finger domain are fused in-frame to the last 191 bp of the UL1 ORF, corresponding to the residues 162 to 224 of gL (Fig. 5a). The splice can be readily detected by end-point RT-PCR using RNA collected from NHDFs infected by multiple HSV-1 strains at 18 hours post infection but not at 6 hours (Fig. 6a). Conversely, the splice is not detected if protein synthesis is blocked using cyclohexamide (CHX) or if viral DNA replication is blocked using phosphonoacetic acid (PAA) or if cells are infected with an ICP4 null mutant defective for early and late gene expression as well as viral origin-dependent replication. Together this suggests that expression of RL2-UL1 requires viral DNA replication. The late kinetics are also evident when the abundance of the RL2-UL1 splice is compared to the canonical RL2 splice by real time RT-qPCR at different times during infection (Fig. 6b). This is notable because ICP0 and gL are considered as immediate-early and late genes respectively. Although canonical UL1 transcripts are not thought to be spliced, the internal UL1 splice acceptor motif GTAG|G is perfectly conserved cross all of the HSV-1 full or partial genome sequences available in GenBank (n = 155) but is not present in HSV-2 genome sequences. Additionally, the RL2-UL1 splice could be detected by RT-PCR in HSV-1 infected ARPE-19 cells and hESC-derived neurons, indicating that this not specific to NHDFs (data not shown).

**Fig. 6:**
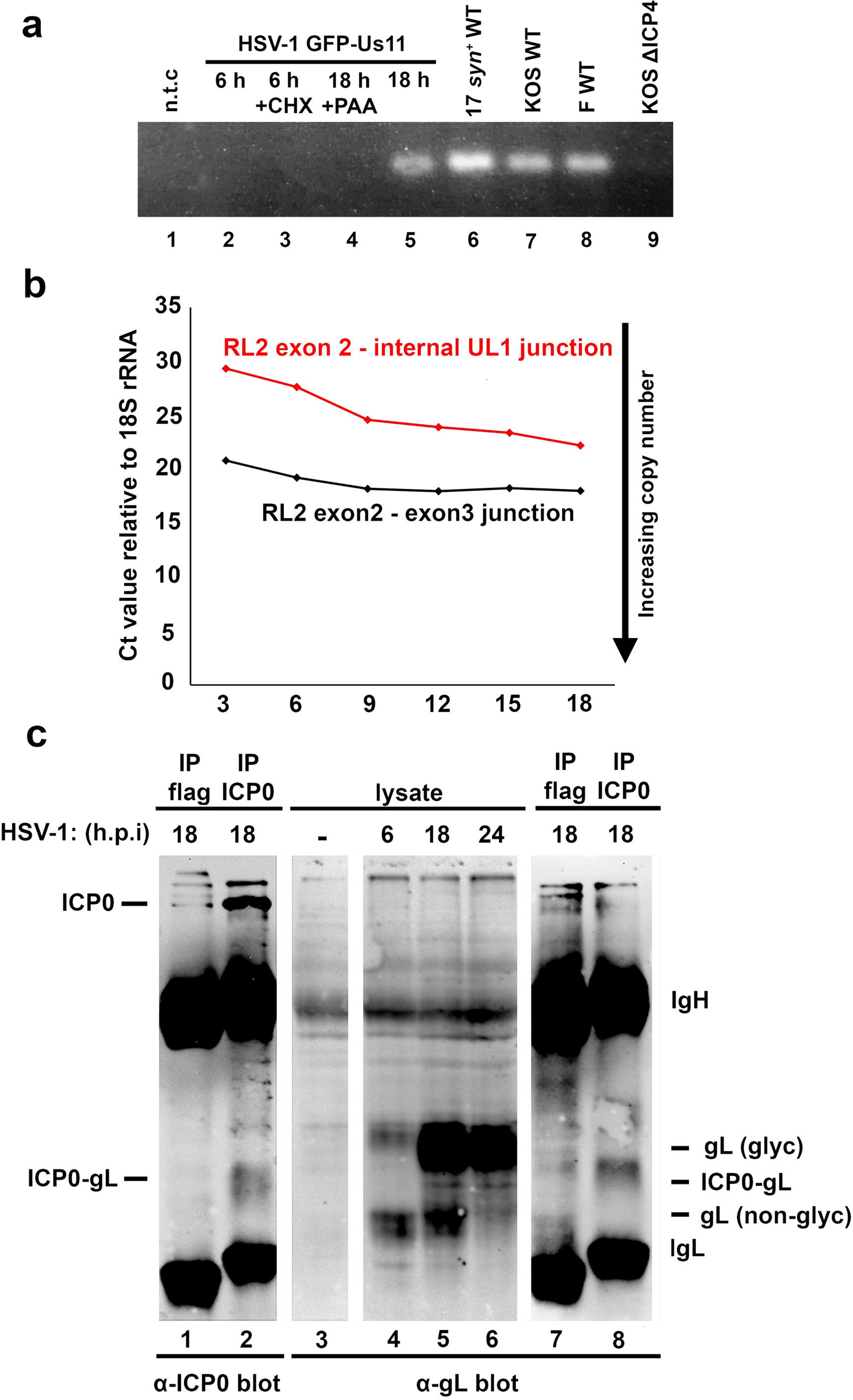
Chimeric RL2-UL1 mRNA is expressed with late kinetics and by multiple HSV-1 strains. **a**, detection of the unique RL2 exon 2 – UL1 splice by RT-qPCR using a primer scanning the splice junction. NHDFs were with infected in parallel with either HSV-1 strain Patton (lanes 2-5) or with wild type strain 17 *syn*+ (lane 6), KOS (lane 7), strain F (lane 8) viruses, or with n12, a KOS ICP4 null-mutant (lane 9), and RNA collected at either 6 h (lanes 2-3) or 18 h (lanes 4-9) post infection. Inhibitors of protein synthesis (cycloheximide, CHX) or the viral DNA polymerase (phosphonoacetic acid, PAA) were included as indicated (lanes 3 and 4). Amplification products were visualized with ethidium bromide. **b**, Assessment of canonical RL2 exon2 - exon3 and novel RL2 exon 2 - UL1 internal splice junction usage at different times after infection by real-time RT-qPCR. Increased RNA abundance is reflected as lower crossover threshold (Ct) values and normalized to 18S rRNA. **c**, Detection of the predicted ICP0-gL fusion protein. Lysates were prepared from mock (lane 3) or HSV-1 strain Patton infected NHDFs collected at 6, 18 or 24 h post infection (lanes 1-2, 4-8) and analyzed by immunoblotting with (lanes 1-2 and 7-8) or without (lanes 3-6) prior immunoprecipitation using anti-ICP0 (lane 2 and 8) or control anti-flag (lanes 1 and 7) loaded protein A beads. After fractionation by SDS-PAGE, membranes were probed using primary antibodies recognizing either the N terminus of ICP0 (lanes 1-2) or the C terminus of glycoprotein L (lanes 3-8). The ICP0-gL fusion peptide has a predicted mass of 32 kDa. Additional reactive species corresponding to ICP0, glycosylated and non-glycosylated gL and antibody heavy (IgH) or light chains (IgL) are indicated.

The predicted ICP0-gL fusion protein has a predicted molecular mass of 32 kDa and a band of this size was detected by immunoblotting with an antibody to the C terminus of gL in fractionated lysates from HSV-1 strain Patton infected NHDFs harvested at 18 or 24 h but not detected at 6 h (Fig. 6c lanes 3-6). To verify this band as the fusion protein rather than a gL glycosylation intermediate, we performed immunoprecipitations on 18 h lysate using either an antibody against the N terminus of ICP0 (lanes 2 and 8) or an irrelevant antibody as a control (1 or 7). Immunoblotting of the recovered material with either α-ICP0 or α-gL detected a diffuse band of similar size in the ICP0-specific immunoprecipitation (lanes 2 and 8) that was absent in the control (lanes 1 and 7). Thus, we are able to confirm expression of this novel HSV-1 gene product, first identified through nRNA-seq, at late times in infection.

## DISCUSSION

Decoding the transcriptional complexity of viral pathogens is vital to understanding how they overcome host cell defense mechanisms and gain control of the host transcriptional and translational machinery. Here, we have systematically examined the utility of native mRNA sequencing using nanopore arrays to profile a highly complex viral transcriptome. We have demonstrated the efficacy and reproducibility of this method and shown that the intrinsic problems associated with the high error rate of nanopore sequencing can be overcome using short-read sequencing data and the generation of what we termed pseudo-transcripts. The resulting transcriptomic data showed greater complexity of viral transcription initiation site usage than previously known and has also revealed an otherwise cryptic class of viral fusion transcripts that encode novel proteins.

Transcription initiation sites (TSS) are critical for productive gene expression as their location relative to the translation initiation site determines the length and composition of the 5′ UTR of mRNAs, which can have profound effects on translation efficiency. Although the majority of RNA polymerase II (RNAP) transcribed genes recruit TFIID to the core promoter, only a minority (~10%) contain an identifiable TATA box^30^. Our data indicates a similar pattern in the HSV-1 genome and in many cases positioning of the TSS may be determined through interactions with ICP4^31^ or through a combination of other core promoter elements^32^. While nRNA-seq does not currently allow mapping of TSS at nucleotide resolution, we nevertheless observed that in many transcripts the 5’ end of HSV-1 sequence reads mapped 30 to 48 bp downstream of a consensus TATA box. This level of resolution is sufficient to identify HSV-1 genes that potentially use multiple TSS, including eight with internalized TSS that are predicted to encode truncated protein isoforms (Supplementary Table 4), four of which have previously been proposed^33^. While beyond the scope of this study, integrating our transciptomic data with alternative approaches such as CAGE-Seq or RNA pol II Chip-Seq should further enhance our understanding of promoter usage in HSV-1 and the relationship of start site selection to subsequent steps in RNA maturation.

Inter-ORF transcription (pervasive transcription) has been observed in several γ-herpesviruses and shown to be functional important^5^. With the exception of polycistronic mRNAs, few examples have been described to date for α-herpesviruses. Detection of HSV-1 fusion transcripts that encode chimeric proteins was therefore exciting and unexpected. Given that the three predominant fusions we identified occur between discrete neighboring transcription units, the likely mechanism involves suppression of transcription termination and polyadenylation leading to elongated transcripts that contain functional splice donor and acceptor sequences. In contrast to our findings, an earlier Illumina-based sequencing study of nascent transcripts in HSV-1 infected fibroblasts found little evidence of read through transcription of viral genes during the first 8 h of infection^18^. This likely reflects the differences in sequencing strategies and analyses methods used. For instance, novel splice site detection is difficult using short Illumina reads and, unlike host transcripts, viral read through transcripts are still able to efficiently terminate because the compact genome provides alternative downstream poly(A) signal sites. While read through transcription remains the most plausible explanation for the generation of fusion transcripts, we cannot entirely exclude the possibility that they instead arise from trans-splicing events that join separate mRNAs. While rare outside of protozoan parasites, there are examples of intergenic splices between viral mRNAs from JC virus and SV40^28, 34, 35^, as well as rarer hybrids between viral and host transcripts.

We focused our subsequent analyses of the predicted ICP0-gL fusion protein encoded by the RL2-UL1 chimeric transcript. While further characterization is required, sensitivity to a viral DNA replication inhibitor and accumulation of the RNA at later times in the infection imply a regulated pattern of expression. Similarly, the conservation of the otherwise unused UL1 splice acceptor sequence amongst all sequenced wild-type HSV-1 genomes, combined with detection of the RL2-UL1 splice in NHDFs infected by four different HSV-1 strains, argues for a functional role for this previously unknown protein. A broader question is whether these viral chimeric mRNAs arise simply a consequence of virus-induced transcription read through targeting the host as part of its host shut off strategy or are primary reason the virus interferes with termination mechanisms.

Technological advances continually redefine our abilities to ask complex questions of biological systems such as the interplay between host and viral transcriptomes during cellular infections. This study illustrates the value of native RNA sequencing and provides a roadmap for researchers interested in examining host-virus interactions that may even be extended to other pathogens (i.e. bacteria, parasites) with transcriptionally complex genomes or loci. The biological insights obtained raises the intriguing question of how many other dsDNA viruses deliberately manipulate the host transcription and RNA processing machinery to increase the diversity of their proteome. Identifying specific mechanisms and determining the role of these mRNA diversification strategies in viral infections will greatly enhance our understanding of host-virus interactions.

### Acknowledgements and funding sources

We thank Hannah Burgess for the R2621 (Δvhs) virus and other members of the Mohr and Wilson Labs, as well as Werner Ouwendijk for useful input. Sequencing was initiated through the NYU Langone Medical Center Genome Technology Center (GTC), which receives support from the National Institutes of Health (NIH) through a grant from the National Center for Advancing Translational Sciences (NCATS UL1 TR00038), and a Cancer Center Support Grant (P30CA016087) to the Laura and Isaac Perlmutter Cancer Center. These studies were also supported individual grants from NIH to IM (AI073898 and GM056927) and ACW (AI130618). T.S. received funding from the Ministry of Education, Culture, Sports, Science and Technology (MEXT KAKENHI JP17H05816, JP16H06429 and JP16K21723), Japan Society for the Promotion of Science (JSPS KAKENHI JP17K008858), the Takeda Science Foundation and Daiichi Sankyo Foundation of Life Science.

### Methods

#### Cell culture, viral strains, and infection procedures

Cells used in this study included normal human dermal fibroblasts (NHDF), human retinal pigment cells (ARPE-19 [ATCC^®^ CRL-2302™]), and hESC-derived neurons^36^. NHDFs were cultured in DMEM supplemented with 5% FBS, ARPE-19s in DMEM/F12 supplemented with 10% FBS and neuronal cultures as previously described^36^. For HSV-1 infections, we utilized multiple viruses corresponding to four different HSV-1 strains including GFP-Us11 Patton^23, 36^, Kos^37^, 17syn^+^, F^38^, KOS N12 (ΔICP4)^39^, and strain F R2621 (Δvhs)^40^, always at a multiplicity of infection (MOI) of three for either 6 or 18 h prior to collecting total RNA or protein. Note that FBS concentrations were halved during the infection and post-infection periods. HSV-1 GFP-Us11 strain Patton is an effectively wild type virus that expresses a fluorescent fusion protein with true late kinetics and has been used extensively in studies of acute infection, latency and viral pathogenesis.

#### RNA collection, extraction, and quality control

HSV-1 infected cells were lysed in TRIzol reagent (Invitrogen) and extracted according to manufacturer’s instructions. RNA integrity (RIN) was assessed using an RNA 6000 nanochip (Agilent Technologies) on a Bioanalyzer 2100 (Agilent Technologies). Poly(A)+ RNA was isolated from 25 μg (HSV -1) or 55 μg (VZV) of total RNA using a Dynabeads™ mRNA Purification Kit (Invitrogen), according to manufacturer’s instructions - with the only adjustment being to use 133 μl resuspended dynabeads rather than 200 μl as this was deemed optimal for the quantity of total RNA.

#### Nanopore sequencing and post-processing

Native RNA sequencing libraries were generated from the isolated poly(A) RNA, spiked with a synthetic Enolase 2 (ENO2) derived calibration strand (a 1.3kb synthetic poly(A)+ RNA, Oxford Nanopore Technologies Ltd.) and sequenced on one of two MinION MkIb with R9.4 flow cells (Oxford Nanopore Technologies Ltd.) and an 18 h runtime. All protocol steps are described in Garalde et al^21^. Following sequencing, base calling was performed using Albacore 1.2.1 [-f FLO-MIN106 -k SQK-RNA001 -r -n 0 -o fastq, fast5]. Only reads present in the ‘pass’ folder were used in subsequent analyses.

#### Illumina sequencing and post-processing

TruSeq stranded RNA libraries were prepared from poly(A)-selected total RNA for the HSV-1 infected NHDFs (18 hpi) and ARPE-19 cells (18 hpi) by staff at the New York University Genome Technology Center. Following multiplexing with additional unrelated samples, paired-end sequencing (2 × 76 cycle) was performed using a HiSeq 4000 (Illumina). FLASh^41^ (--min-overlap=10 --max-overlap=150) was used to merge overlapping reads prior to error-correction.

#### Error correction of native RNA nanopore reads

For error-correction of nanopore sequence reads, we utilized proovread^26^, an error-correction package initially designed for data correction for the PacBio long-read sequencing platform. Here, we generated error-corrected nanopore datasets by applying proovread to nanopore sequence reads using subsampled FLASh^41^ compacted Illumina RNA-Seq datasets (250,000 – 5,000,000 paired-end reads) generated from the same material. As a metric we examined per-read changes in CIGAR string lengths, reasoning that correction of indels and substitution-type errors would reduce string lengths and improve splice-site usage identification (Supplementary Fig. 2). Interestingly, inspection of individual error-corrected nanopore reads showed no clear pattern in terms of how many subsampled Illumina reads were optimal. We therefore utilized a nomination protocol (Fig. 2b) to identify the best corrected version of each read across all datasets (scored by shortest CIGAR string length) and generated a final error-corrected dataset to take forward into the next step of our analysis (Fig. 2a). Error-correction conveyed two main benefits. The first was to increase the number of reads mapping to the HSV-1 genome/transcriptome while the second was to improve minimap2 alignments around splice sites (Fig. 1 and 2).

#### Generation of pseudo-transcripts

While error-correction significantly reduced the number of indel-type errors in our nanopore reads, it rarely removed all of them. Thus attempts to identify and translate ORFs within these reads produced frameshift errors that obscured the likely ORF sequence. However, by leveraging the start, stop and internal splicing site co-ordinates from the mapping output, we were able to generate what we term ‘pseudo-transcripts’ by substituting in the corresponding reference genome sequence in place of the existing nanopore read sequence (Fig. 2a). This served to repair sequences to an extent that internal ORF predictions could be made. Note that while error-correction is not strictly required for the generation of pseudo-transcripts, the improved mapping accuracy, particularly of splice junctions, justifies the additional computational time required.

#### Mapping of nanopore sequence data

Following basecalling with Albacore, nanopore reads are separated into three folders (pass, fail, calibration strand). We used only the reads in the pass folder for subsequent analyses and for each dataset, we ran the analyses with both raw (uncorrected) and proovread-corrected datasets. Nanopore read data were aligned to the HSV-1 strain 17A reference genome (NC_001806) and transcriptome, as well as the *Homo sapiens* HG19 genome and transcriptome using MiniMap2^24^ (-ax splice -k14 -uf --secondary=no), a splice aware aligner. Due to many HSV-1 genes being arrayed as polycistrons, we utilized two distinct version of the transcriptome, one containing all encoded ORFs (used for internal splicing analysis and isoform clustering), the other containing all encoded transcriptional units (utilized for examining transcript boundaries and fusion transcript discovery). Note that following mapping, all SAM files were parsed to sorted BAM files using SAMtools v1.3.1^42^.

#### Data Visualization

Figures showing read data overlaying genome schematics were generated in RStudio (http://www.rstudio.com) using GViz and GenomicFeatures packages^43, 44^.

#### Transcript boundaries analysis

BAM files containing read data mapped to the HSV-1 genome were parsed to BED12 files using BEDtools^45^, separated by strand, and the extreme 5’ mapping (*cut -f2 BED12 | sort -n | uniq -c | sed ′s/^ *//′ | sed ′s/ /\t/g′ | awk ′{print "HSV1-st17",$2,$2,$1}′*) and 3’ mapping (*cut –f3 BED12 | sort -n | uniq -c | sed ′s/^ *//′ | sed ′s/ /\t/g′ | awk ′{print "HSV1-st17",$2,$2,$1}′*) sites identified for all mapping reads. The resultant dataset is a four column file specifying each unique 5’ start site and 3’ end site identified and the number of distinct transcripts utilizing that start site. These can be visualized as described above.

#### Fusion transcript analysis

BAM files containing read data mapped the HSV-1 transcriptome (transcriptional units) was parsed to identify putative chimaeras (SAM flag “SA:Z”). Each chimeric sequence read was translated to identify all ORFs >30 amino acids using ORFfinder (https://www.ncbi.nlm.nih.gov/orffinder/). The resulting peptide sequences were queried (blastp^46^) against a blast database comprising all canonical HSV-1 proteins. The resulting data was manually parsed to identify peptide sequences mapping to two or more ORFs present in distinct transcriptional units. Subsequent visualization using IGV^27^ was used to identify splice acceptor and donor sequences and to verify mapping integrity.

#### Examination of RL2-UL1 splice site usage

Primers were designed to discriminate between usage of the canonical RL2 exon2 – exon3 splice junction and the novel RL2 exon – UL1 internal splice junction (Fig. 6, Supplementary Table 6). Primers designed against the *H. sapiens* 18srRNA were utilized as a control. cDNA was generated from 500 ng total RNA using qScript cDNA SuperMix XLT (Quanta Bio) and 50ng cDNA used per PCR / real-time quantitative PCR (qPCR) reaction. For PCR amplification, the Herculase II Fusion DNA Polymerase (Agilent Technologies) was used according to manufacturer’s instructions in a 25 μl reaction volume and with 35 amplification cycles. PCR products were visualized alongside a GeneRuler 100 bp ladder (Thermo Scientific) on a 1.8% agarose gel stained with ethidium bromide. For Real time qPCR, all reactions were carried out in triplicate using a reaction volume of 25 μl and the SsoAdvanced™ Universal SYBR^®^ Green Supermix (BIO-RAD). All samples were run on a CFX96-Touch (Bio-Rad).

#### Immunoprecipitation and immunoblotting

Cells were lysed in 1x cell lysis buffer (Cell Signaling), fractionated by 15% sodium dodecyl sulfate polyacrylamide gel electrophoresis (SDS-PAGE), and transferred to nitrocellulose membranes (Whatman). Membranes were blocked in 5% milk diluted in Tris Buffered Saline with Tween^®^ 20 (TBS-T) for 1 h at room temperature and incubated overnight at 4°C with α-ICP0 (1:200, 53070, Santa Cruz Biotechnology), or α-gL^47^ (1:1000, H1A259-100, Virusys Corp) primary antibodies and detected using a horseradish peroxidase-conjugated α-mouse secondary antibody (diluted 1:5000, Sigma-Aldrich) with incubation at room temperature for 1 h and visualization by chemiluminescent detection using SuperSignal™ West Femto Maximum Sensitivity Substrate (ThermoFisher). For immunoprecipitation, lysates were incubated for 1 h at 4°C with α-ICP0 or α-FLAG loaded protein G agarose beads (Cell Signaling), recovered by low speed centrifugation, and washed in TBS before heat denaturation in sample buffer. Images were captured using an iBright FL1000 (Invitrogen).

#### Data availability

Basecalled nanopore and Illumina fastq datasets generated as part of this study can be downloaded from the European Nucleotide Archive (ENA) under the following study accession: PRJEB27861. Raw fast5 files are freely available from authors upon request.

---

**Supplementary Fig. 1:**
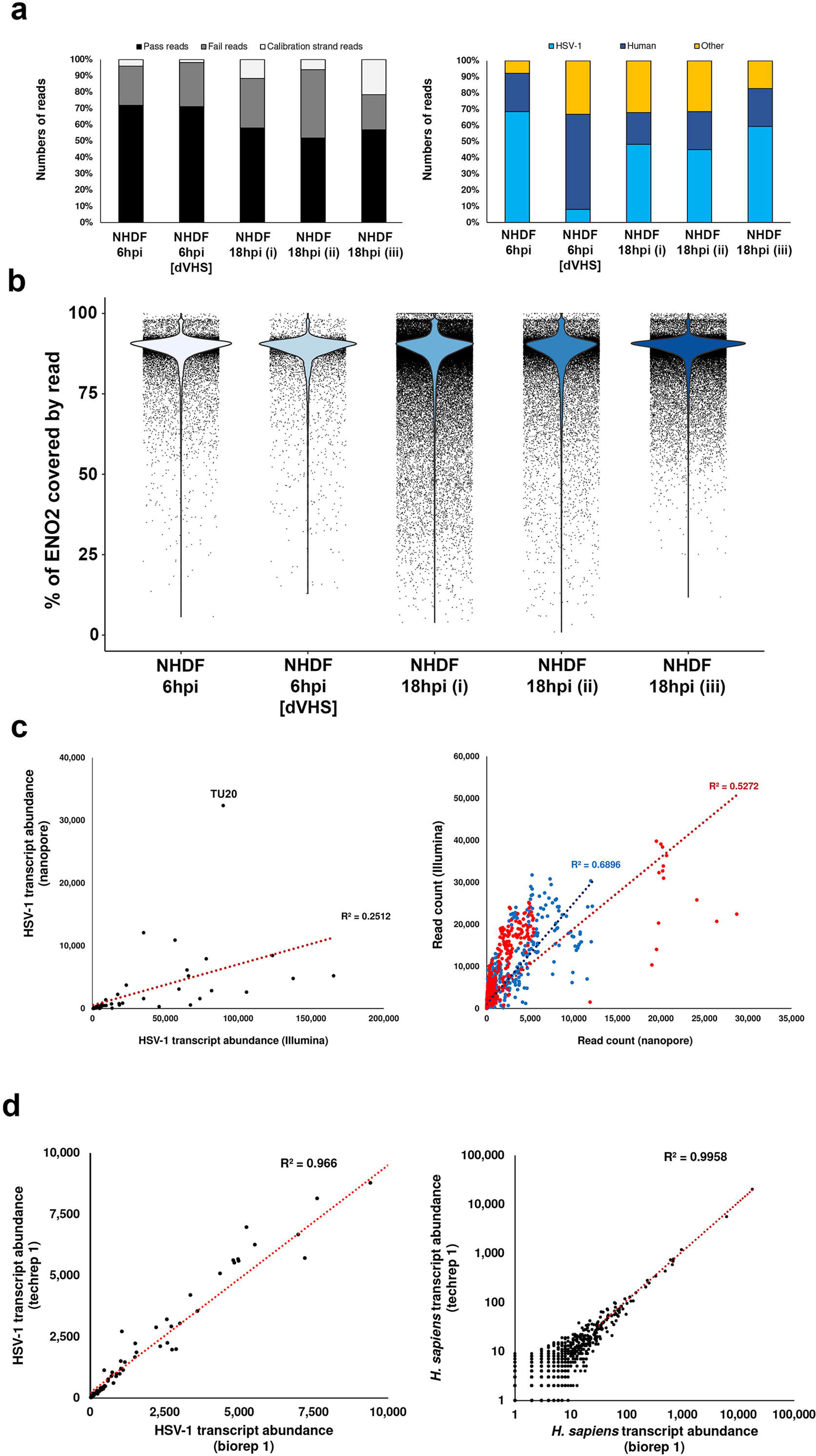
mRNA is minimally degraded during library construction and sequencing. **a**, Alternative summary of quality control (left) and genome origin (right) metrics for five separate native RNA sequencing runs from normal human dermal fibroblasts (NHDF) infected with either HSV-1 strain Patton GFP-Us11 or HSV-1 strain F vhs null virus (dVHS) and the RNA harvested after 6 or 18 h. **b**, the spiking of ENO2 mRNA allows assessment of mRNA degradation during library preparation. Here, mRNA degradation is represented by the fraction of ENO2 covered by individual reads and indicates only minimal 5’ degradation during library preparation. **c**, Correlation analyses of HSV-1 transcript abundances (left) and HSV-1 genome coverage (right) generated using nanopore and Illumina sequence data. The sliding window analysis is generated by calculating and plotting mean read depth values per 100 nucleotide windows in a strand specific manner (red - top strand, blue - bottom strand). **d**, transcript abundances were counted for the HSV-1 (left) and *H. sapiens* (right) transcriptomes and showed near perfect correlation between technical replicates of the NHDF 18hpi (i) and NHDF 18hpi (iii) samples, sequenced on separate MinION devices.

**Supplementary Fig. 2:**
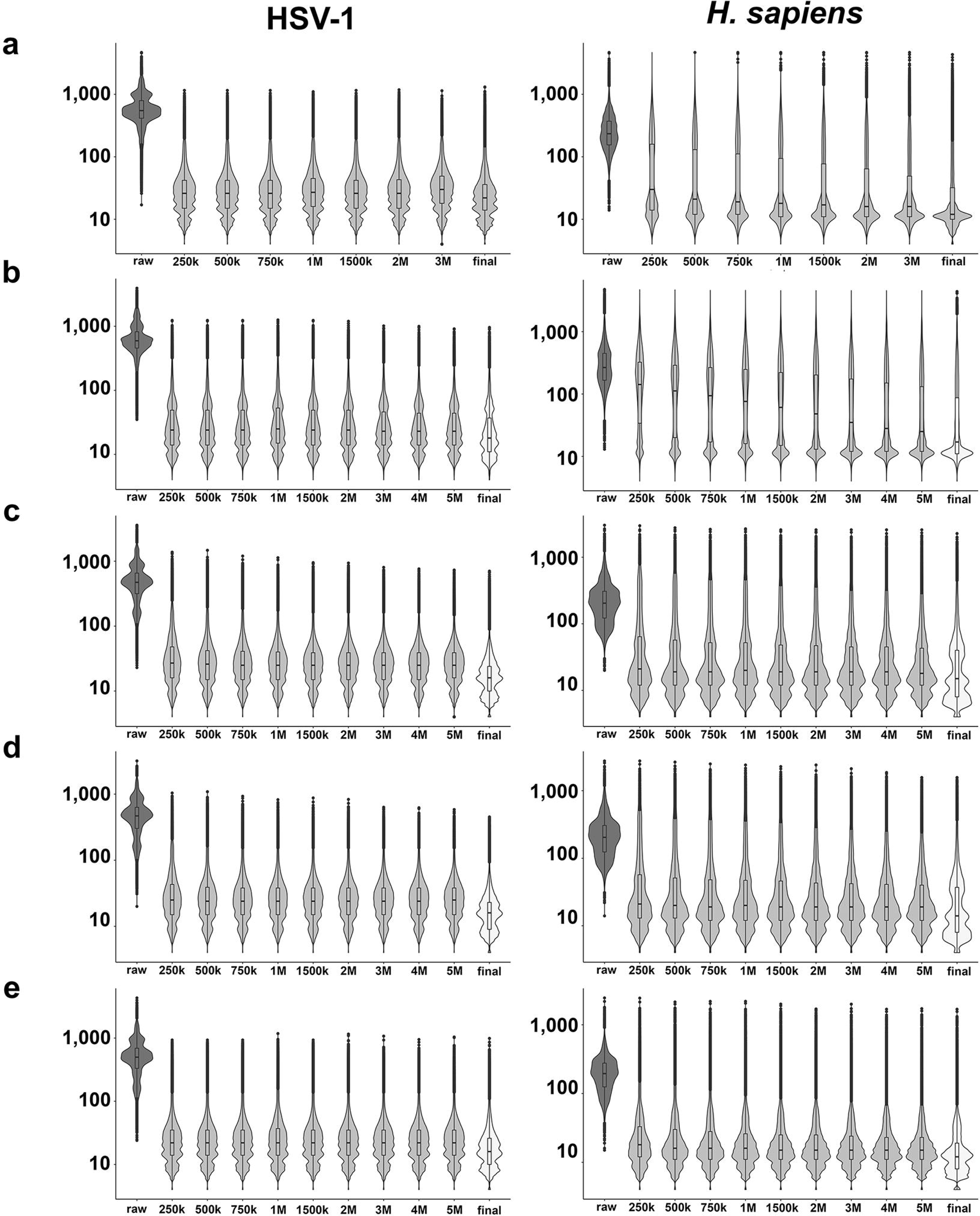
Proovread error-correction significantly reduces the impact of indel and substitution type errors in nanopore direct RNA reads. For each of the five datasets generated in this study, sequence reads were mapped against the HSV-1 strain 17 (left) and *H. sapiens* (right) genomes and the CIGAR string length (y-axis) extracted for each individual read. Here, violin plots show the distributions of CIGAR string lengths and the effect of error-correction using proovread and FLASh-merged Illumina sequencing libraries, subsampled for varying numbers of reads (x-axis). The raw dataset comprises uncorrected nanopore reads while the final dataset contains the optimally corrected reads derived from a decision matrix (Fig. 2b), scored by shortest CIGAR string length. **a**, NHDF 6 hpi **b**, NHDF 6 hpi [HSV-1 dVHS mutant] (ii) **c**, NHDF 18hpi (i) biological replicate **d**, NHDF 18hpi (ii) biological replicate and **e**, NHDF 18hpi (iii) technical replicate on separate minION device.

**Supplementary Fig. 3:**
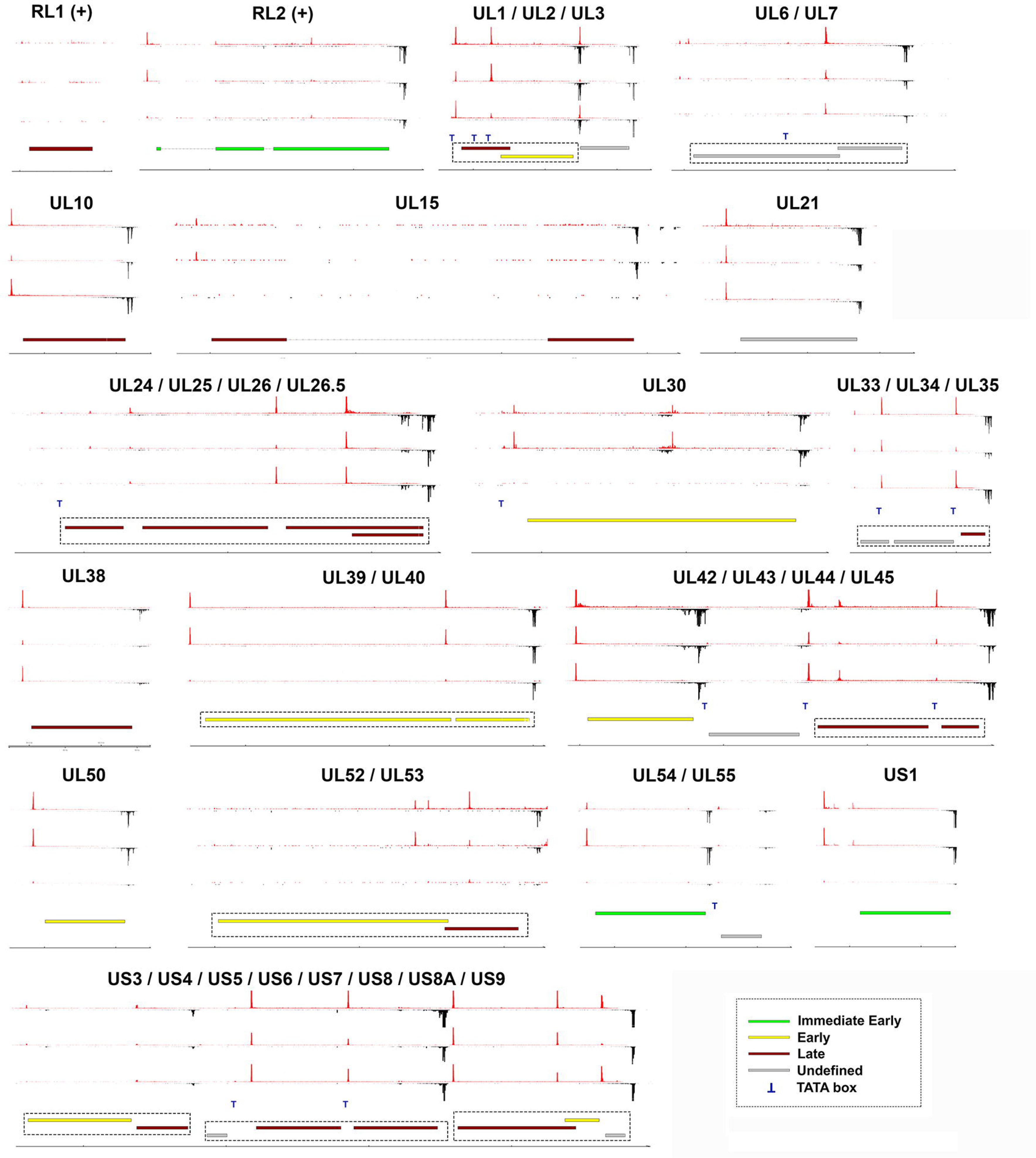
Identification of proximal transcription initiation and termination sites along the top strand of the HSV-1 genome. Data sets correspond to NHDF infected with HSV-1 strain Patton for 6 hpi (upper track), strain F dVHS for 6 hpi (middle track) and strain Patton for 18 hpi (lower track). Transcripts are initiated a few nucleotides from the extreme 5’ end of each nRNA-seq read (red peaks) and polyadenylation (black peaks) occurs downstream of canonical (AAUAAA) poly(A) signal sequences. Canonical HSV-1 ORFs are colored according to kinetic class (IE – green, E – yellow, L – red, undefined – grey) while polycistronic transcriptional units are indicated by hatched boxes. Identifiable TATA boxes are indicated by a blue T. Transcription proceeds from left to right. Note only the regions encompassing canonical ORFs are shown.

**Supplementary Fig. 4:**
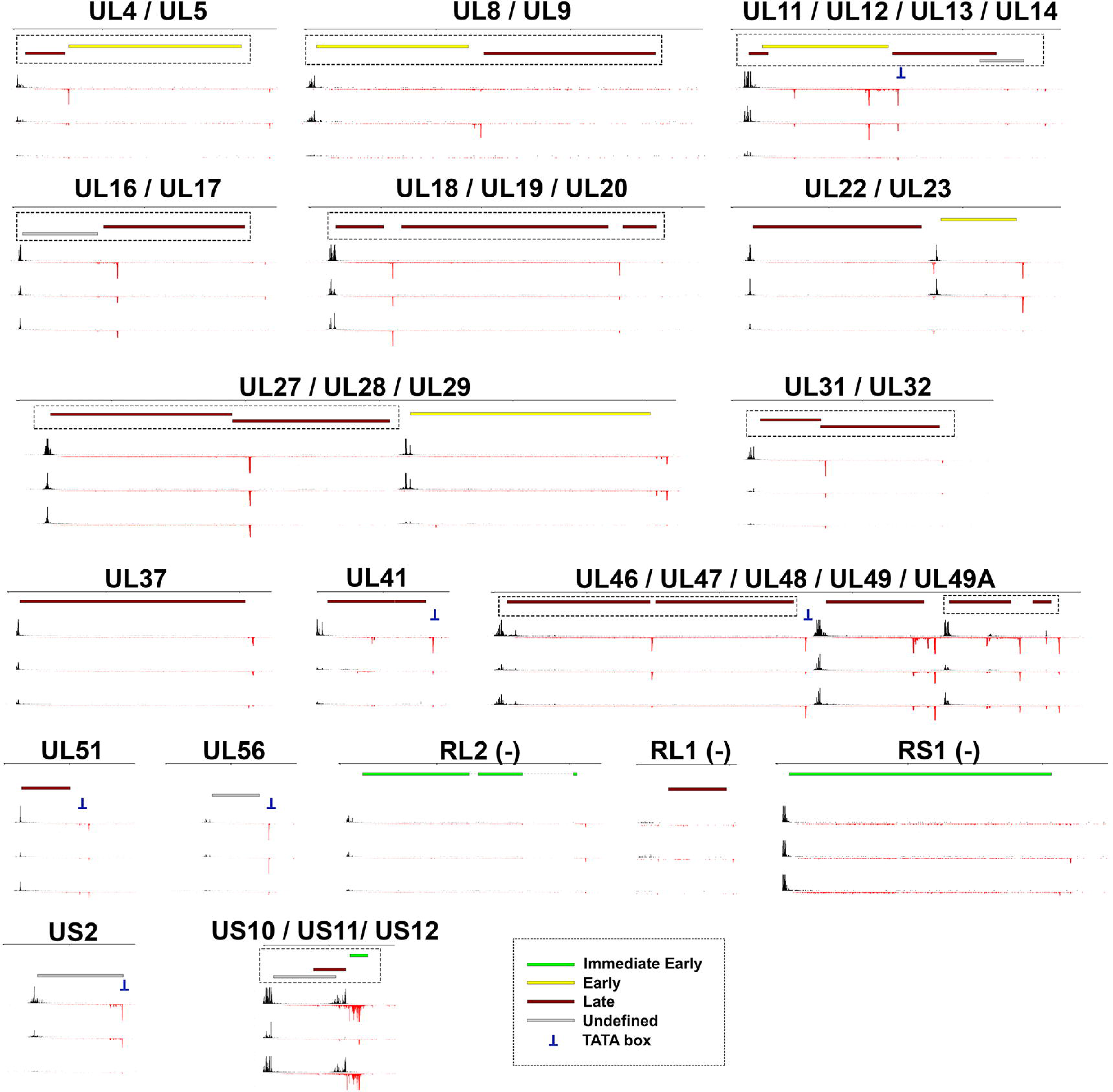
Identification of proximal transcription initiation and termination sites along the bottom strand of the HSV-1 genome. Data sets correspond to NHDF infected with HSV-1 strain Patton for 6 hpi (upper track), strain F dVHS for 6 hpi (middle track) and strain Patton for 18 hpi (lower track). Transcripts are initiated a few nucleotides from the extreme 5’ end of each nRNA-seq read (red peaks) and polyadenylation (black peaks) occurs downstream of canonical (AAUAAA) poly(A) signal sequences. Canonical HSV-1 ORFs are colored according to kinetic class (IE – green, E – yellow, L – red, undefined – grey) while polycistronic transcriptional units are indicated by hatched boxes. Identifiable TATA boxes are indicated by a blue T. Transcription proceeds from right to left. Note only the regions encompassing canonical ORFs are shown.

